# Using social network analysis and non-invasive antibody detection to explore pathogen exposure in wildlife communities

**DOI:** 10.64898/2025.12.09.693199

**Authors:** Oriane Ploquin, Oriane Basso, Mthabisi Ndlovu, Franck Prugnolle, Alexandre Caron, Martin Munzamba, Emildah Porovha, Khanyile Nkomo, Gaëlle Corbel, Céline Arnathau, Anne Boissière, Valérie Pinarello, Nina Rault, Richard Shumba, Masotsha Mhlanga, Tawanda Tarakini, Simon Chamaillé-Jammes, Hervé Fritz, Florian Liégeois, Ellen Mwandirigana, Constance Sibelo, Chaitezvi Columbus, Hillary Madzikanda, Vladimir Grosbois, Eve Miguel

## Abstract

Understanding the dynamics of infectious diseases in ecosystems shared with wildlife is a priority for preventing future threats to human, animal health and conservation. However, the collection of ecological and epidemiological data on wild populations has relied on invasive and costly methods which limited the capacity of investigation. Recent technological developments have changed this trend. In Hwange national park and its surrounding area in Zimbabwe, we combined camera trap-based ecological monitoring over a period of 12 months, covering 14 water holes with a fecal-based antibody survey in 16 large herbivore species. A survey involving 52,829 pictures as well as 629 faecal samples collected every 15 days, has shown Through the modelling of multispecies contact networks, the risk of infection exposure at species-level is predicted. Coupled with foot-and-mouth disease virus (FMDV) antibody information, the role of each species in the dynamics of infectious diseases is explored. This study highlights how community networks can provide valuable insights into the functional epidemiological role of wildlife populations. Rather than establishing transmission routes, our aim is to propose a scalable and non-invasive surveillance framework that identifies priority species and areas for epidemiological monitoring in complex ecological systems.

**Significance Statement:** Monitoring disease circulation in wildlife is often hindered by the difficulty of collecting epidemiological data. We propose a novel non-invasive approach that combines species interaction networks derived from camera-trap data with non-invasive antibody detection to explore the exposure patterns of large herbivore communities to foot-and-mouth disease virus (FMDV). By linking the position of species in contact networks with their immunological status, we demonstrate the potential of using ecological centrality as a proxy for identifying key hosts in transmission or indicator species for pathogen circulation. Beyond FMDV, this framework can be adapted to other pathogens for which non-invasive immunological assays are currently being developed, making it broadly relevant for wildlife disease surveillance in remote or protected areas.

## Introduction

More than half of human diseases are of animal origin (1, 2). Infectious diseases such as Ebola, Zika, Nipah, and avian influenza demonstrate the intrinsic link between human, animal and ecosystem health. Preventing future crises relies on a comprehensive approach to global health (3).

Among the many components shaping savanna ecosystem health, large herbivores of the orders Artiodactyla and Perissodactyla play a particular role. They share numerous pathogens with humans (4) and, through the long history of livestock expansion, have been partly domesticated, often living in close proximity to humans (5). While studies on the circulation of diseases in livestock are common (6–8), the epidemiology of infectious diseases in wild ungulates remains largely understudied (9, 10).

Individual-level infection data, although not essential in all epidemiological contexts,including modeling, remains a critical resource for capturing pathogen dynamics (Fitzner & Heckinger, 2010). Collecting wildlife samples to produce epidemiological data typically involves invasive methods such as captures and sedations (12, 13). Recently, these approaches are increasingly being avoided, not only because of the logistical challenges they pose, but also due to growing concerns raised by animal welfare considerations and advocacy from animal rights groups. Developing alternative methods is a priority, as this would significantly improve our understanding of wildlife health within the context of public health and conservation issues (14).

Foot-and-mouth disease (FMD) is a multi-host pathogen that infects a wide range of domestic and wild ungulates. Despite decades of research, its transmission dynamics remain poorly understood, which continues to hinder effective control. The high morbidity in livestock and the movement restrictions imposed as control measures are the main drivers of its substantial economic impact in affected regions (15). Globally, FMD is estimated to circulate in 77% of livestock populations (16). Beyond livestock, the disease also affects wildlife populations, complicating conservation efforts and transboundary management (17). Wildlife and livestock can both act as maintenance hosts depending on the epidemiological context, further increasing the complexity of pathogen circulation as the number of host species rises (4, 18).

The first step in addressing the functional role of hosts and non-hosts in disease transmission is to describe the social structures of at-risk wildlife communities (19, 20). This functional approach categorizes species based on their epidemiological roles: maintenance hosts are capable of sustaining pathogen circulation independently, bridge hosts facilitate transmission between maintenance hosts and other susceptible species, and dead-end hosts become infected but do not contribute to further transmission (18, 21). Identifying these roles allows targeted assessment of which species are likely to drive, mediate, or limit disease spread. Maintenance populations have the capacity to sustain the pathogen over the long-term without external inputs; they function in the multi-host system as a reservoir for the pathogen.

By grouping populations based on the link between epidemiological functions and ecological traits, we can develop eco-epidemiological models to manage transmission dynamics of infectious diseases transmission across diverse species (18, 21).

In multi-species systems, it is crucial to identify the key hosts that play a vital role in the amplification of pathogens within communities. These superspreader populations form strong connections within the network, centralising their role in the community’s interaction dynamics and contributing significantly to pathogen circulation as efficient hosts. This contrasts with the role of non-competent host species, which act as dead ends for virus transmission due to their inefficiency in spreading the virus. These highly connected hosts can be characterised through social network analysis using metrics such as node centrality (20, 22–24).

By linking the characterisation of contacts between functional groups with epidemiological data, we can determine the role of each species within a multi-host epidemiological system. This provides undeniable added value to disease management strategies (25). Although existing eco-epidemiological studies often monitor the serological incidence of a single species or family as an indicator of infectious dynamics within the community (26, 27), achieving a comprehensive understanding of the system remains elusive. This is because surveying multiple populations simultaneously is challenging, despite this being crucial for a holistic view of infectious dynamics.

Based on the position of species within social networks, we propose to use faecal samples as a promising substrate for detecting proxy of infection (i.e virus or antibodies). Fecal samples are reliable substrates for the detection of immunoglobulin A (IgA) antibodies (28), which are exclusively secreted by mucosal surfaces (29). These antibodies have shown significant potential for enhancing our understanding of infectious diseases, particularly with regard to identifying persistent infections in carriers (30), or simply determining the infection status/pathogen exposure of an animal (29, 31). Biologists already use faecal samples in studies to assess dietary niche partitioning (32), for example, to investigate steroid secretion as a stress response (33) or for pathogen surveillance (34). There are numerous advantages to use faecal samples over serum samples, with the primary benefit being the ease with which a large number of samples can be collected non-invasively.

This study presents the results of a one-year ecological and epidemiological survey of a sub-Saharan wildlife community comprising sixteen species of large ungulates that are either hosts or non-hosts of the foot-and-mouth disease (FMD) virus. We demonstrate the effectiveness of a new, non-invasive, integrated monitoring approach. This method reliably addresses the complexity of multi-host infectious disease systems and provides new insights into identifying maintenance, superspreader and indicator species in the spread of FMD.

Taken together, our study provides a scalable, non-invasive framework for understanding pathogen dynamics across communities, offering new opportunities to integrate wildlife into global disease surveillance.

## Results

### Ecological centrality varies widely across species in wildlife contact networks

Over the course of one year, camera traps deployed at 15 water points resulting in 52,829 images capturing the target species, from which 83 interspecific contact networks were constructed based on the compilation of data per site and per season. These contact networks reflect the structure of wildlife interactions at a community level, based on temporal co-occurrence at water sources. Using seroprevalence reported in literature, transmission networks have been modeled on the basis of contact networks to picture the contribution of each species in the circulation of FMD virus within each of the 83 sessions (Figure 1, table 1).

**Figure 1.**
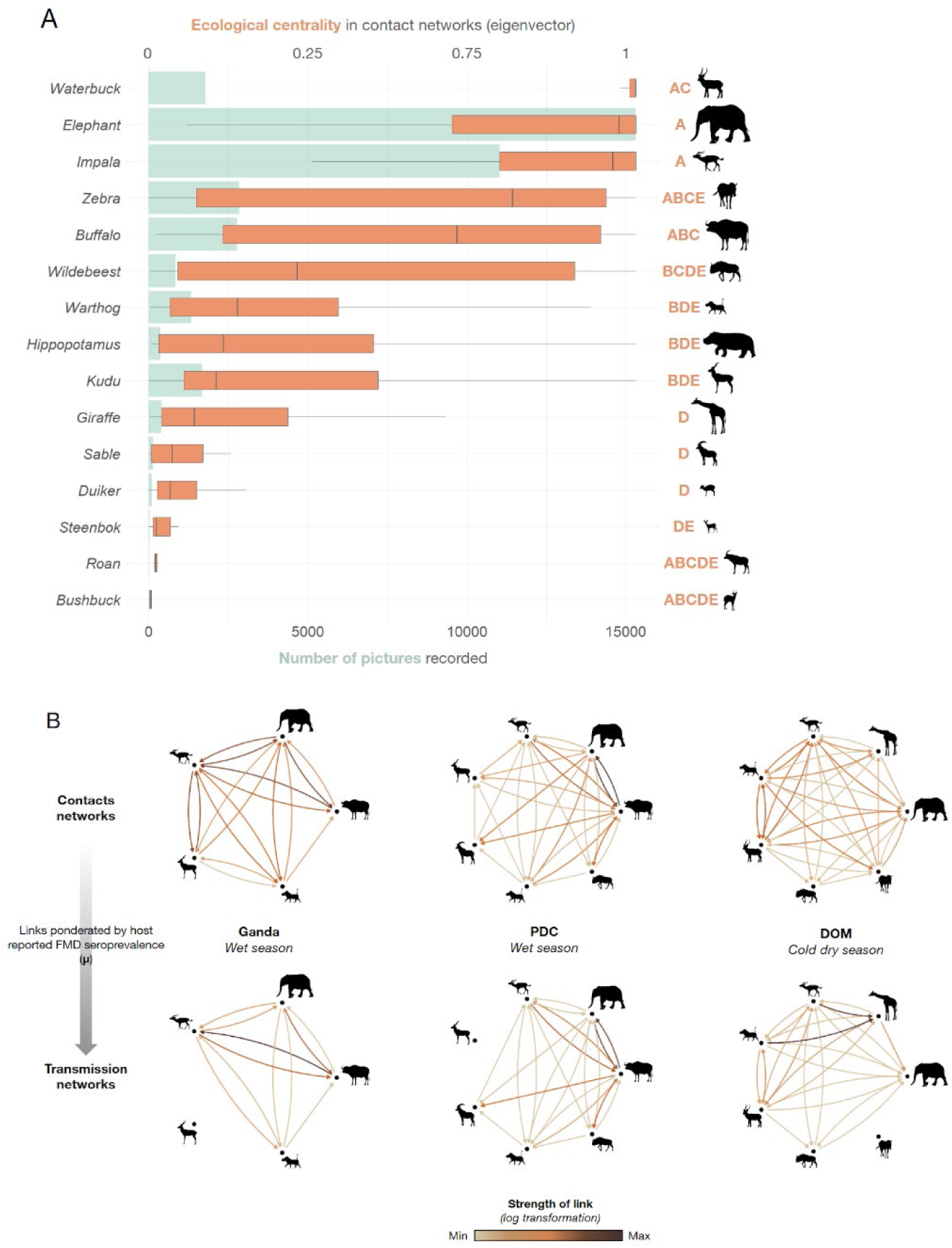
(A) Detection and centrality of the species captured during the camera trap based survey. The species studied are ranked by median eigenvector centrality (in orange), a proxy of abundance of the species is provided through the number of pictures taken for each species (in green). Different letters represent significant differences among species according to Wilcoxon tests. (B) Weighted and directed contact networks built based on the camera-trap-based survey (first line), links are weighted by the seroprevalence of both species included in the link reported in the literature to obtain transmission networks. Darkness of the links are positively associated with the strength of contacts between two species.

**Table 1.**
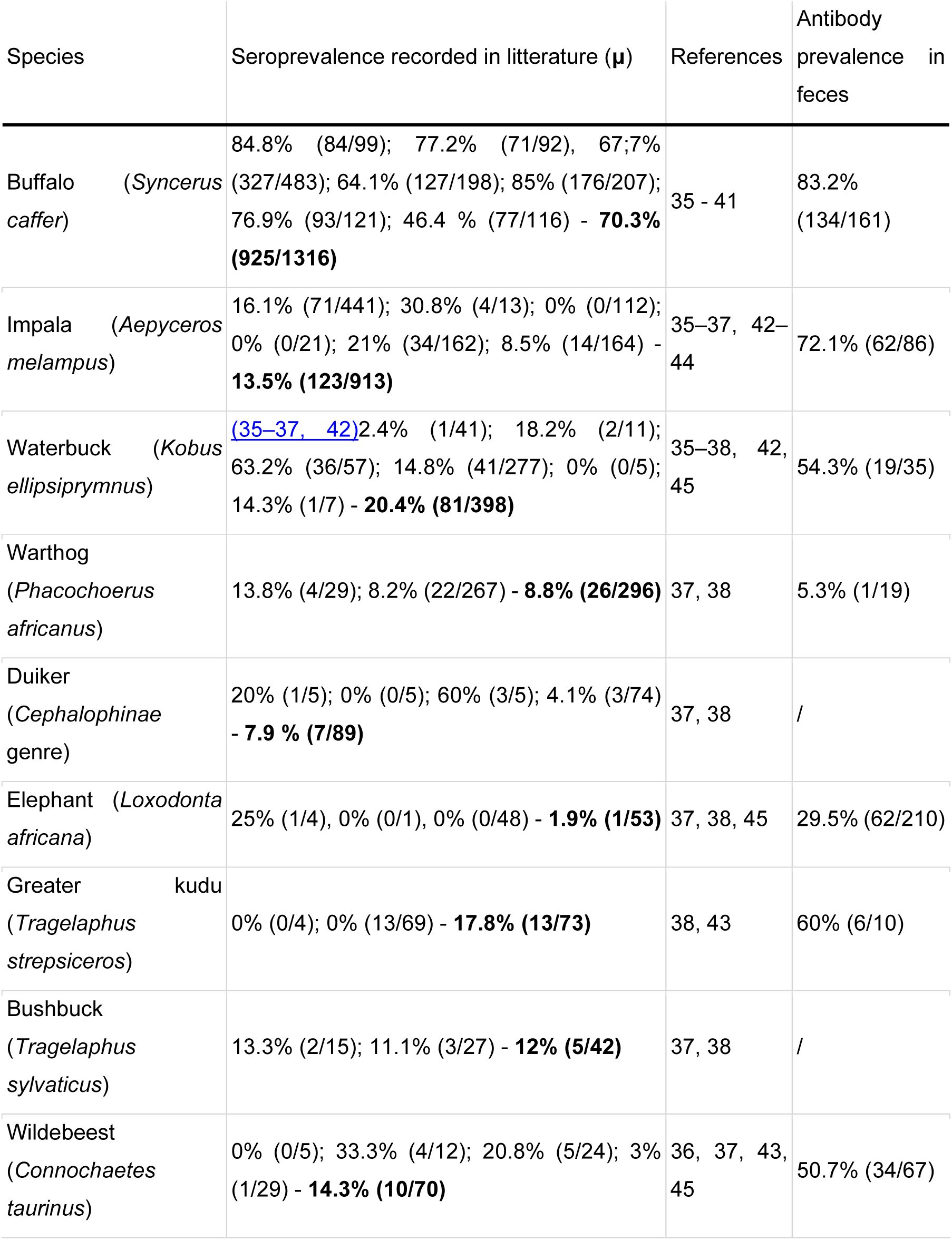
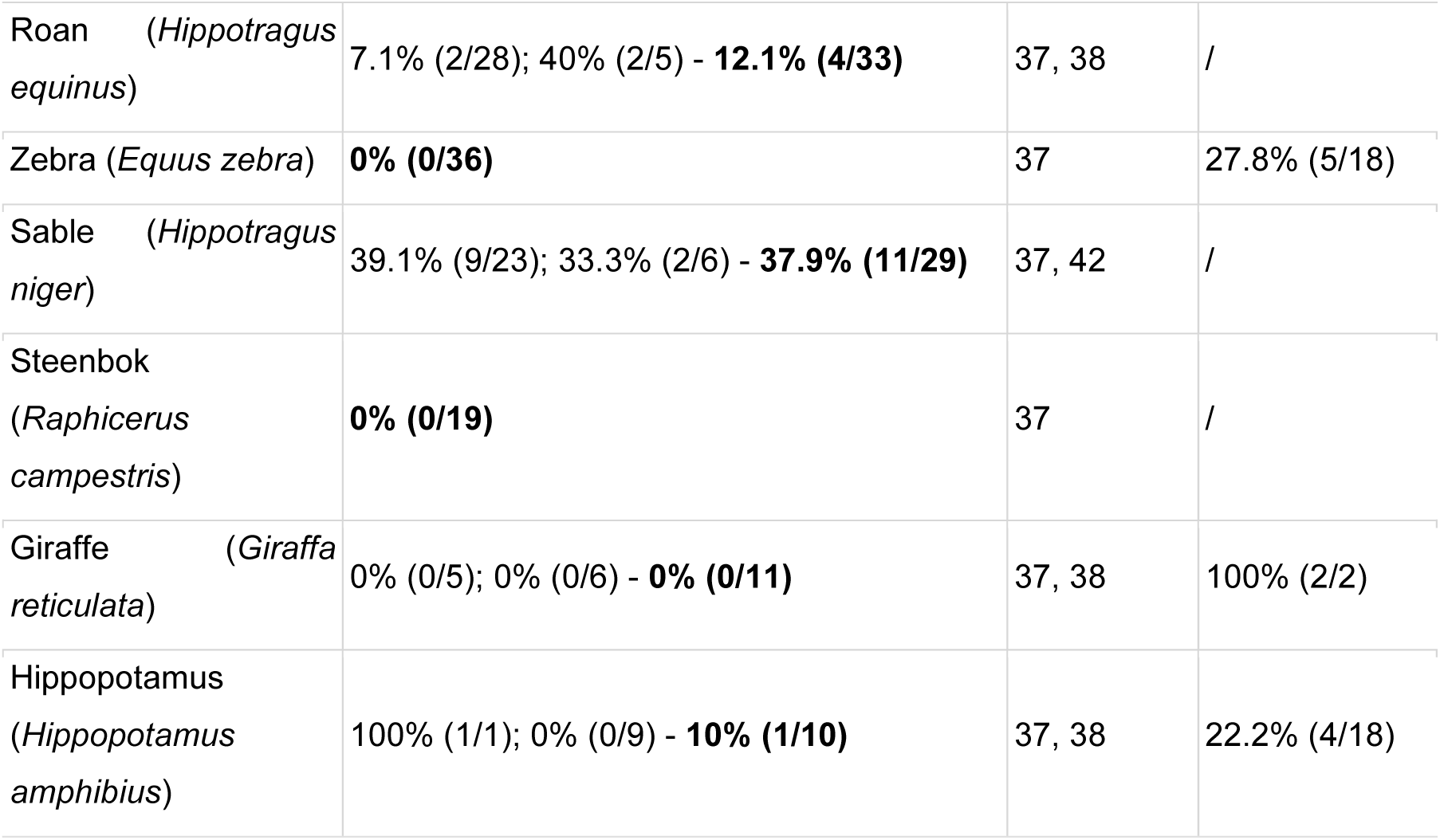
Prevalence of FMD-specific antibodies reported in the literature and detected in fecal samples for this study.

Among the 15 herbivore species detected, elephants, impalas, zebras, and buffaloes were the most frequently observed. Network analysis revealed significant differences in species ecological centrality across the community. Eigenvector centrality, a metric capturing both direct and indirect connections, varied markedly among species in the contact networks (Friedman test, χ² = 891.01, df = 14, p < 2.2 × 10⁻¹⁶, Figure 1).

Impalas, waterbucks, elephants, and wildebeests consistently exhibited the highest centrality scores, indicating strong positions in the interaction network. In contrast, giraffes, duikers, and bushbucks occupied relatively more peripheral positions.

These results suggest marked variation in how species mediate interspecific contact within the community. However, network centrality does not equate to species abundance, as the metric is independent of individual counts.

### Fecal antibody detection reflects known seroprevalence patterns

A total of 626 fecal samples were collected across 10 focal species around water holes. FMDV-specific antibodies were detected in several species using a commercial ELISA targeting non-structural proteins (NSP). Despite the exploratory nature of fecal antibody detection, the prevalence estimates aligned closely with published serological data. To contextualize our findings, we systematically compiled literature estimates of FMD seroprevalence obtained from wild populations of the same subspecies present in our study area. We restricted our review to studies based on serum samples and excluded domestic or captive populations, but applied no specific geographic or temporal restrictions. This ensured comparability while maximizing the available evidence. References to the studies used for comparison are provided in Table 1. A linear regression weighted by the number of samples per species showed a significant positive relationship between seroprevalence reported in the literature and prevalence observed in feces (slope = 0.724, SE = 0.167, *p* = 0.003), with an intercept of 34.89 (SE = 6.27, *p* < 0.001). The model explained 69.7% of the variance (R² = 0.697), indicating that reported seroprevalence is a strong predictor of fecal prevalence, though some variability remains. A leave-one-species-out sensitivity analysis indicated that the relationship was largely driven by buffalo, whose exclusion substantially reduced the strength and significance of the association (see SI Appendix). Nonetheless, the effect size remained positive, suggesting that species with higher network centrality tended to exhibit higher antibody prevalence overall.

This concordance suggests that fecal samples may contain immunological signals indicative of prior pathogen exposure at the population level, despite not being validated as diagnostic material (Table 1, Figure 2).

**Figure 2.**
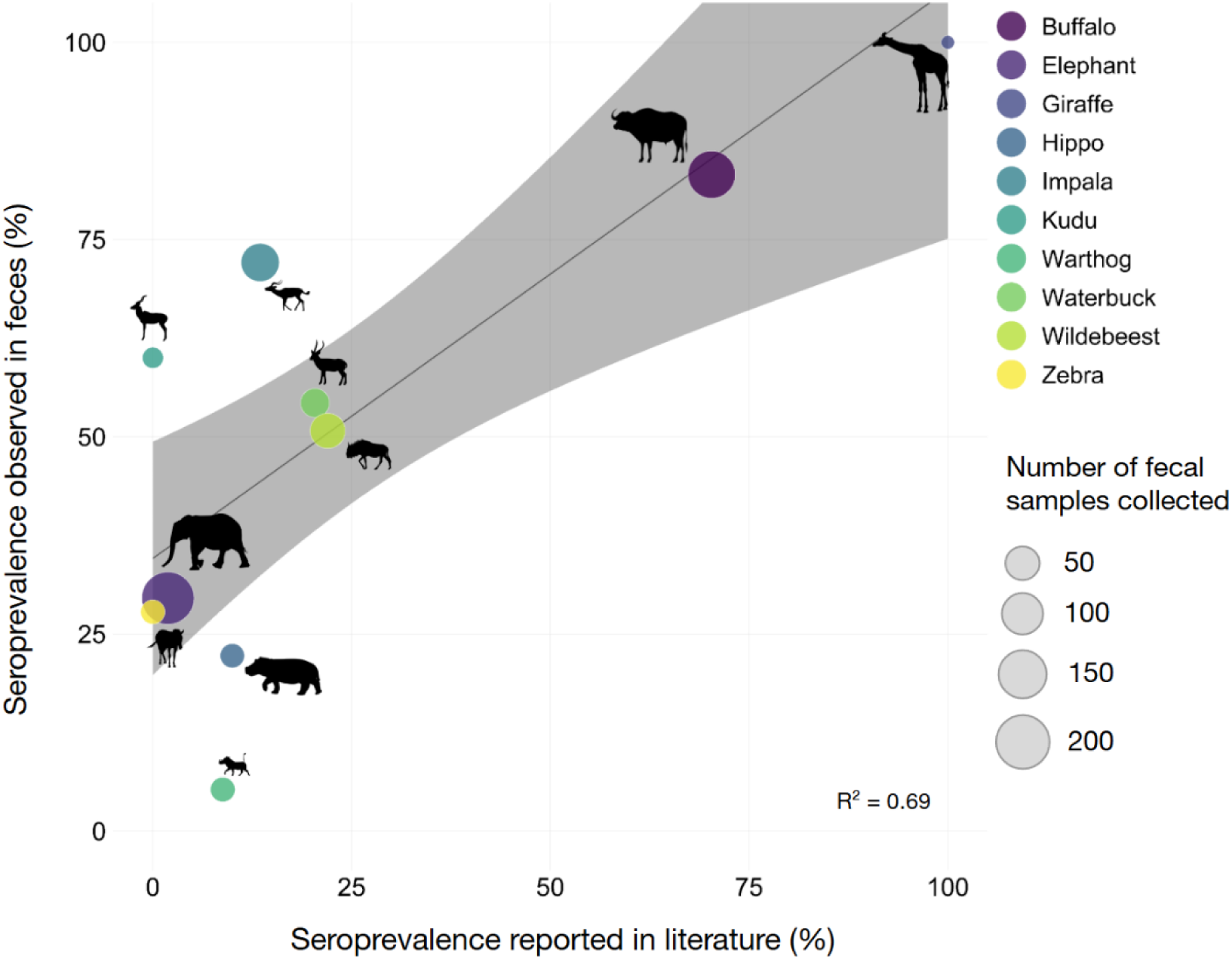
Weighted linear regression between the seroprevalence of foot-and-mouth disease virus (FMDV) antibodies reported in the literature and the prevalence observed in fecal samples. Each point represents one host species, with color indicating species identity and point size proportional to the number of fecal samples analyzed. The regression line (in grey) is weighted by sample size.The sampling size of the literature can be assessed in Table 2.

**Table 2.** Fixed effects of the GLMM predicting FMD seropositivity effects of the GLMM predicting FMD seropositivity.

| Fixed effects | Estimate | Std. Error | z value | p value |
| --- | --- | --- | --- | --- |
| (Intercept) | -1.22 | 0.45 | -2.68 | 0.007** |
| Exposure to pathogen (pagerank) | 418.79 | 91.17 | 4.59 | < 0.001*** |
| Ecological centrality (eigenvector) | 0.16 | 0.45 | 0.36 | 0.7254 |

Significance codes: 0 \*\*\*, 0.001 \*\*, 0.01 \*.

### FMDV antibodies detection in feces increases with species’ exposure in the modelled transmission network

To explore whether contact network position predicts pathogen exposure, we transformed the contact networks into potential transmission networks. Links between species (i.e., nodes) were weighted by both the frequency of their contacts and estimated host competence (derived from literature-based seroprevalence, μ, Table 1 & Figure 1), yielding a directed network reflecting plausible transmission pathways.

Using these transmission networks, we computed the absolute PageRank (number and strength of direct and upstreaming links received by a node) as a proxy for species exposure to the virus. A generalized linear mixed model including site and season as random intercepts explained 27% of the total variance in FMD seropositivity in feces (conditional R² = 0.27; marginal R² = 0.12). PageRank centrality had a strong positive effect on the probability of detecting FMD antibodies (β = 418.8 ± 91.2, z = 4.59, p < 0.001), whereas eigenvector centrality showed no significant association (p = 0.72). Random effects indicated moderate variability among sites (SD = 0.78) and lower variability among sampling seasons (SD = 0.34)(Table 2, Figure 3).

**Figure 3.**
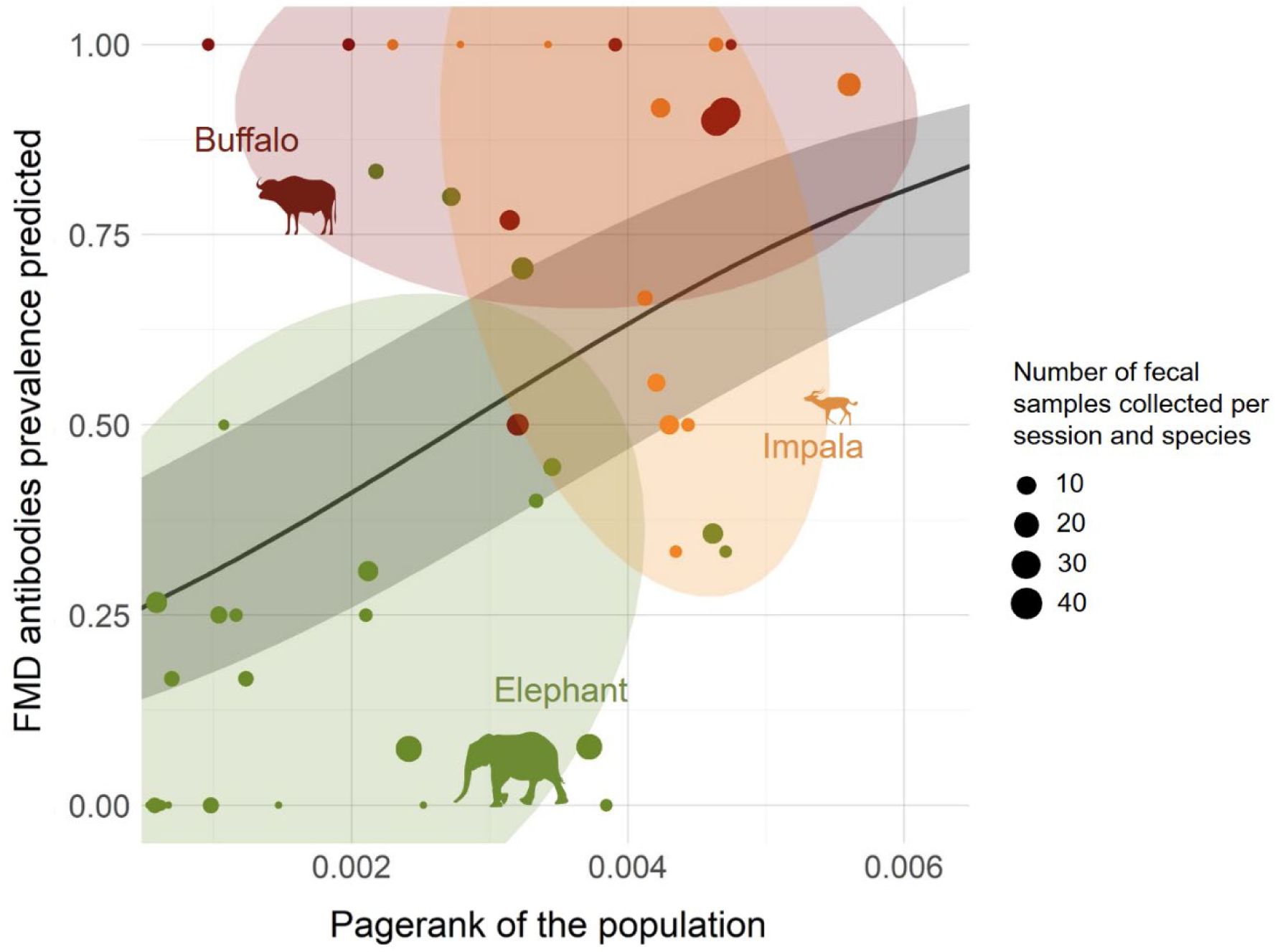
Relationship between species’ exposure in the modelled transmission network and Foot-and-Mouth Disease Virus (FMDV) antibody prevalence in wild ungulates. The grey line and shaded ribbon represent the fitted values and 95% confidence interval predicted by a generalized linear mixed model (GLMM; binomial family, logit link) including species’ PageRank (reflecting both the number and strength of incoming transmission links) as a fixed effect, and site and season as random intercepts. Colored points and ellipses depict empirical data: each point shows the mean observed FMDV antibody prevalence per species and sampling unit (point size proportional to sample size), and ellipses represent the 80% confidence region for each species. Colors indicate species—buffalo (dark red), impala (orange), and elephant (olive green).

## Discussion

Ecosystem-wide active epidemiological surveillance is essential to understanding disease risks at the wildlife–livestock–human interface. In systems where animal-related industries such as tourism and livestock are central to the economy, spillover events pose serious threats to both biodiversity and public health. Monitoring such risks requires scalable approaches that can operate across species and landscapes, while minimizing disturbance to wildlife populations.

This study introduces a methodological framework that integrates ecological contact networks using camera traps with non-invasive antibody detection in feces to investigate pathogen exposure in a diverse wildlife community. While exploratory and limited by the absence of assay validation for fecal matrices, the approach revealed patterns broadly consistent with established seroprevalence data—particularly driven by buffalo—suggesting that fecal antibodies may capture meaningful immunological signals, at least for species with high exposure levels. Rather than providing a diagnostic tool, this proof of concept demonstrates the value of combining ecological and immunological data to generate testable hypotheses about host roles in pathogen circulation. Estimates of host competence, though derived from heterogeneous serological studies, nonetheless yielded consistent cross-species patterns, underscoring the potential of this integrative framework despite methodological constraints. However, challenges with this approach include the presence of inhibitors and a lack of understanding of antibody production profiles against foot-and-mouth disease (FMD) in different populations, such as IgA, IgD, IgE, IgG and IgM, which are produced at different stages of infection, such as initial exposure, or in response to different types of infection, such as allergic or parasitic (46).

By coupling this immunological signal with ecological network metrics, we identified species likely to play structurally important roles in pathogen exposure. Specifically, impalas and elephants emerged as highly central nodes in our contact networks, frequenting water points and interacting widely with other species. These ecological patterns align with what is known about FMD ecology: impalas are recognized bridge hosts, capable of linking maintenance (buffalo) and recipient (cattle) populations (47, 48), while elephants—despite similar connectivity—are not competent for FMD transmission, making them potential ecological dead-ends. This illustrates how ecological centrality and epidemiological function may differ across pathogens, but also how centrality can serve as a screening metric to identify species worth monitoring in multi-host systems. Central species like elephants, although not competent transmitters and therefore only exposed to the virus through other species, may nonetheless act as early indicators of community-level circulation or as buffers that modulate transmission flow. Conversely, highly central and competent hosts, such as impalas, may play disproportionate roles in amplification or spillover.

We also show that network-derived exposure scores (PageRank) are significantly associated with the presence of antibodies in fecal samples. This relationship supports the idea that ecological contact structure can be used to infer exposure risk, even in the absence of individual-level tracking or invasive sampling. In this sense, our framework contributes to the development of eco-epidemiological models that integrate behavioral, environmental, and immunological data.

Importantly, our framework is inherently scalable. The use of camera traps and fecal samples allows for repeated, cross-sectional surveys that are both low-cost and non-invasive. Moreover, variations in antibody prevalence between sites—despite potential overlaps in animal movement—suggest that our method can detect localized immunological dynamics, possibly reflecting spatial heterogeneity in pathogen circulation.

While our findings focus on FMD, the approach can be extended to other pathogens for which antibody detection in feces is feasible or under development. The framework also opens opportunities to couple environmental DNA, microbiome profiles, or molecular detection of pathogen genomes with behavioral data to refine exposure models. For example, elephants are known transmitters of pathogens such as anthrax and bovine tuberculosis (49, 50). Depending on the pathogen, these networks could help clarify how a single species may shift between distinct epidemiological roles across disease systems.

In summary, we propose a novel, non-invasive approach to explore pathogen exposure in wildlife communities. By integrating species interaction networks with exploratory fecal immunology, we identify structurally important species and ecological contexts associated with increased exposure. Although further validation is needed, particularly on the immunological side, our results provide a strong rationale for incorporating network structure into wildlife disease surveillance strategies.

This work contributes to bridging a critical gap between ecological monitoring and pathogen risk assessment in free-ranging animal communities. In doing so, it supports broader efforts to integrate biodiversity conservation, animal health, and public health within a unified One Health framework.

## Materials and Methods

### Study system

The study was conducted in a protected savanna ecosystem comprising the Sikumi Forest Area (1,100 km²) and adjacent zones at the eastern periphery of Hwange National Park, Zimbabwe (26°9’E, 18°6’S). This area, part of the Kavango-Zambezi Transfrontier Conservation Area, hosts one of the largest populations of free-ranging African elephants, alongside a diverse community of large herbivores including buffalo, impala, kudu, roan, zebra, and others (51). The region receives ∼600 mm of annual rainfall and lacks perennial water bodies, relying on seasonal or pumped waterholes that serve as ecological hubs (52).

### Camera trap deployment and species detection

Fourteen water points were randomly selected and monitored using two camera traps per site (Browning Dark Ops HD Max), positioned approximately 10 m from the water’s edge at a height of 1.5 m. Cameras operated for 15 days per month between June 2022 and July 2023, using 1-minute time-lapse during the day and motion-triggered recording at night. The initial dataset comprised over 3,300,000 images. Empty images were first removed by MegaDetector algorithm provided by WildEye, and the remaining images were processed using the open-source software TrapTagger also provided by WildEye (https://wildeyeconservation.org/), which performed a first automated species classification. All identifications were subsequently verified manually. Only large herbivore species relevant to FMD ecology (53) or locally abundant were retained for analysis (16 species in total).

### Contact network construction

Contact events were defined as detections of two different species at the same water point within a 5-day window, consistent with estimates of FMDV persistence in the environment (54). Contact strengths were aggregated into weighted, directed networks across 15-day sessions and sites. All analyses were performed in R (version 3.6.0+; R Core Team, 2025) using RStudio (version 2025.09.0+387; RStudio Team, 2025). All monitored species for both camera-traps deployment and feces collection, as well as literature sources reporting FMDV seroprevalence, are listed in Table 1.

### Transmission network modeling

Contact networks were transformed into transmission networks by weighting links according to species-level transmission potential (µ), approximated by average seroprevalence from published data. Centrality metrics included: Eigenvector centrality (to identify potential bridge/superspreader species based on ecological position in the contact network), Absolute PageRank (to estimate cumulative exposure risk in the transmission network), normalized across all networks (Table 1, see SI Appendix).

Network metrics were computed using igraph (55) and sna (56).

### Fecal sampling and antibody detection

A total of 626 fresh fecal samples (<24h) were collected at water points during the camera-trap survey period every 15 days. Samples were identified to species by experienced field personnel and stored in Viral Transport Medium (VTM) consisting of 100 ml phosphate-buffered saline (PBS), 1 ml penicillin-streptomycin solution, and 0.05 ml (one drop) amphotericin B (mycostatin) at −20°C until processed. Detection of antibodies against FMDV non-structural proteins (NSP) was performed using a commercial ELISA kit (ID Screen® FMD NSP Competition, ID.vet, France). Fecal supernatants were obtained by double centrifugation and assayed according to manufacturer instructions. Detailed procedures are described in the Appendix.

This assay was not originally developed for fecal matrices; results were interpreted as exploratory proxies for exposure, not diagnostics.

### Statistical analysis

Species differences in network centrality metrics (eigenvector centrality and PageRank) were assessed using Friedman tests, followed by pairwise Wilcoxon rank-sum tests. The relationship between fecal antibody prevalence and literature-based seroprevalence was evaluated using weighted linear regression. We tested the effect of species PageRank on antibody prevalence using a Generalized Linear Mixed Model (GLMM; lme4 package)(49), including site as a random effect.

## Acknowledgments

This work was supported by the HUM-ANI project of the Climate and Biodiversity Initiative, funded by the BNP Paribas Foundation; the BIO-PATHO project (ECOPATHO, CNRS Sciences Frugales 2021); and the WHISHES exploratory project of LabEx CeMEB (Centre Méditerranéen Environnement et Biodiversité, Montpellier University). We thank the Zimbabwe Parks and Wildlife Management Authority and the Forestry Commission of Zimbabwe for granting us access to their land and facilitating the collection of data with the support or field rangers. We are also grateful to the safari operators who facilitated our work: Amalinda CoSafari Collection (Khulu, Ivory, Sable Valley), Hwange Safari Lodge, Ganda Lodge, Iganyana Lodge, and Sikumi Tree Lodge. We acknowledge our partners for assistance with camera-trap image analysis: TrapTagger, WildEye, and Innoventix Consulting (Pty). Research permits were granted by ZimParks (23(1)(C)(II)11/2022 and 23(1)(C)(II)15/2023).

## Notes

### Competing Interest Statement

The authors have declared no competing interest.

