## Supporting information for "Using social network analysis and non-invasive antibody detection to explore pathogen exposure in wildlife communities"

---

### Calculation of contact strength

Contacts between species were inferred from co-occurrence events at water points within a temporal window of ≤5 days. The strength of contact $C_{AB}$ between species A and B was calculated using an exponential decay function based on the time lag $\Delta t$ between detections:

$$C_{AB} = \sum_{i=1}^{n} e^{-\lambda \Delta t_i}$$

Where:

- A and B are two species (nodes) in the contact network
- $\Delta t_i$ is the time difference in days between detections in event $i_1$
- $\lambda = \log(2)/2.5$ corresponding to a half-life of 2.5 days

For each 15-day session and site, these values were aggregated to generate a weighted, directed adjacency matrix.

---

### Transmission potential weighting

Transmission potential between species was modeled using:

$$T_{A \to B} = C_{AB} \times \mu_A \times \mu_B$$

Where:

- A and B are two species (nodes) in the transmission network
- $\mu_A$ and $\mu_B$ are species-level proxy values of transmission competence (based on compiled seroprevalence data; Table 1)
- $C_{A \to B}$ is the normalized contact strength.

This generated a matrix used for computing absolute PageRank centrality.

---

### Networks metrics used for this study

We used two complementary centrality metrics to characterize each species' position within the interspecific transmission network: **Eigenvector centrality** and **PageRank**. Both quantify how connected a node is within the network, but they differ in how they weight direct and indirect connections.

**Eigenvector centrality**

Eigenvector centrality measures the influence of a node in the network by considering both the number and the quality of its connections. A node connected to many highly connected neighbors will have a high eigenvector score, while a node connected to poorly connected nodes will have a lower score. Mathematically, the eigenvector centrality $x_i$ of node *i* is defined as:

$$x_i = \frac{1}{\lambda}\sum_{j=1}^{n} A_{ij}x_j$$

where $A_{ij}$ is the element of the adjacency matrix representing the connection between nodes *i* and *j*, and $\lambda$ is the largest eigenvalue of *A*. This recursive definition implies that a node's centrality depends not only on its direct links but also on the centralities of its neighbors. Eigenvector centrality thus captures the global influence of a node within the entire network.

**PageRank**

PageRank centrality extends the logic of eigenvector centrality by incorporating the direction and weighting of links, as well as a *damping factor* that accounts for the probability of following a connection in a stochastic process. Originally developed by Google to rank web pages, PageRank estimates the likelihood that a random walker moving along the network will arrive at a given node. Formally, the PageRank of node *i* is computed as:

$$PR(i) = \frac{1-d}{N} + d \sum_{j \in M(i)} \frac{PR(j)}{L(j)}$$

where $d$ is the damping factor (typically 0.85), $N$ is the total number of nodes, $M(i)$ is the set of nodes linking to $i$, and $L(j)$ is the number of outgoing links from node *j*. PageRank therefore reflects both the number and the strength of incoming connections, as well as their importance within the broader network. In our context, PageRank represents the cumulative probability of receiving direct or upstream transmissions, and can thus be interpreted as a proxy for exposure risk to the virus.

**Fecal Sample Processing for Serological Analysis**

***Sample Thawing and Preparation***

Fecal samples were removed from the –20 °C freezer the evening before processing and placed at 4 °C overnight to allow gradual thawing. Pasteur pipettes were prepared by cutting the tip to enlarge the opening, taking care not to make it too wide, which could hinder accurate aspiration.

***Sample Processing and Decontamination***

The thawed fecal suspension was centrifuged for 10 min at 4,500 rpm, ensuring that tubes were balanced in the centrifuge. The supernatant was carefully aspirated using a Pasteur pipette and transferred into a 15 mL tube, which was centrifuged again for 10 min at 4,500 rpm. Approximately 1 mL of the resulting supernatant was then transferred into a 1.5 mL cryotube. Sample tubes were placed in a water bath at 56 °C for 2 hours for decontamination.

***Fecal Sample Preparation for Serology***

Decontaminated fecal matter was aspirated using a prepared Pasteur pipette and transferred into a 5 mL tube. Tubes were vortexed to homogenize the sample and centrifuged for 10 min at 13,000 rpm, ensuring proper balance in the centrifuge and inward orientation of caps. After centrifugation, the supernatant was separated from the pellet and immediately used in a competitive ELISA for the detection of Foot and Mouth Disease 3ABC nonstructural protein antibodies (NSP) in serum and plasma from bovine, ovine, caprine, porcine and all susceptible species (ID Screen® FMD NSP Competition).

---

**Sensitivity analysis: leave-one-species-out**

To assess the robustness of the relationship between observed FMD seroprevalence (mean_elisa_fmd) and compiled seroprevalence (seroprevalence), we performed a leave-one-species-out (LOO) analysis. For each iteration, one species was removed from the dataset and a weighted linear model was fitted:

mean_elisa_fmd ~ seroprevalence, weights = count

The slope estimate, standard error (SE), $R^2$, and p-value were recorded for each iteration (Table S1).

**Table S1.** Leave-one-species-out sensitivity analysis

| Species removed | Estimate | SE | R_squared | P_value |
|---|---|---|---|---|
| Buffalo | 1.144 | 0.567 | 0.37 | 0.083 |
| Elephant | 0.577 | 0.205 | 0.53 | 0.026 |
| Giraffe | 0.723 | 0.185 | 0.69 | 0.00578* |
| Hippo | 0.710 | 0.178 | 0.70 | 0.00521* |
| Impala | 0.787 | 0.103 | 0.89 | 0.000126** |
| Kudu | 0.734 | 0.178 | 0.71 | 0.00448* |
| Warthog | 0.698 | 0.162 | 0.73 | 0.00356* |
| Waterbuck | 0.723 | 0.182 | 0.69 | 0.00539* |
| Wildebeest | 0.721 | 0.183 | 0.69 | 0.00556* |
| Zebra | 0.714 | 0.184 | 0.68 | 0.00604** |

*Significance codes: 0 ***, 0.001 **, 0.01 *.*

*Note:* Estimates correspond to the slope of the linear model relating observed vs. compiled FMD seroprevalence, weighted by species-level sample counts. This analysis tests the sensitivity of the relationship to the removal of individual species.

**Model comparison for GLMMs**

We tested four candidate models with varying fixed and random structures. The final model included:

- Response variable: antibody presence (binomial),
- Fixed effect: transmission PageRank, eigenvector centrality
- Random effect: water point ID. season (wet, cold dry, hot dry)
